# PhysMAP - interpretable *in vivo* neuronal cell type identification using multi-modal analysis of electrophysiological data

**DOI:** 10.1101/2024.02.28.582461

**Authors:** Eric Kenji Lee, Asım Emre Gül, Greggory Heller, Anna Lakunina, Santiago Jaramillo, Pawel F. Przytycki, Chandramouli Chandrasekaran

## Abstract

Cells of different types perform diverse computations and coordinate their activity during sensation, perception, and action. While electrophysiological approaches can measure the activity of many neurons simultaneously, assigning cell type labels to these neurons is an open problem. Here, we develop PhysMAP, a framework that weighs multiple electrophysiological modalities simultaneously in an unsupervised manner and obtain an interpretable representation that separates neurons by cell type. PhysMAP is superior to any single electrophysiological modality in identifying neuronal cell types such as excitatory pyramidal, PV^+^ interneurons, and SOM^+^ interneurons with high confidence in both juxtacellular and extracellular recordings and from multiple areas of the mouse brain. PhysMAP built on ground truth data can be used for classifying cell types in new and existing electrophysiological datasets, and thus facilitate simultaneous assessment of the coordinated dynamics of multiple neuronal cell types during behavior.

## Introduction

Single cell transcriptomics, *in vivo* and *in vitro* electrophysiology, and morphological reconstruction have identified numerous neuronal cell types in the cerebral cortex (Scala et al., 2021; Gouwens et al., 2020;, BICCN) each with specific patterns of gene expression, morphology, and connectivity (Bomkamp et al., 2019). These interconnected cell types form cortical microcircuits which perform neural computations and, ultimately, produce behavior (Keller et al., 2020; Kullander and Topolnik, 2021; Christensen et al., 2022). For instance, recent studies have identified how interactions between excitatory pyramidal, parvalbumin-positive (PV^+^), and somatostatin-positive (SOM^+^) cells enable gain modulation (Ferguson and Cardin, 2020). These interactions have been used to model computation for functions such as the selective enhancement of certain visual stimuli (Millman & Ocker et al. 2020).

However, how individual cell types contribute to the many other computations the brain conducts is largely unknown (Adesnik and Naka, 2018), as direct measurement of specific types necessitates molecular techniques (e.g., optogenetics and calcium imaging) that are not feasible in all situations. Such techniques are far from turnkey and are not easy to execute due to various factors e.g., a lack of viral serotype efficacy (Bohlen and Tremblay, 2023), the need for optical access and head-fixation (O’Shea et al., 2017), or the depth of relevant neural populations (Trautmann et al., 2021) etc. Even when these techniques are available, technical constraints restrict experimentation: only a few cell types are imageable at once and two-photon imaging is limited to cortical depths of around 600 *µ*m (Takasaki et al., 2020). This precludes the study of cell types from deep structures and in many of the species most relevant for understanding human disorders such as non-human primates (Bliss-Moreau et al., 2022). Furthermore, in order to develop precision circuit-level therapies, cell type-specific activity is needed to validate microcircuit models as network properties are predictive of therapeutic response (Bloch et al., 2022). Thus, our current inability to capture cell type dynamics hinders our understanding of how circuits orchestrate behavior in intact and diseased states.

In contrast to optical methods, extracellular electrophysiological recordings from single electrodes to high-density Neuropixels probes (Jun et al., 2017) can sample from diverse and deep populations of neurons simultaneously and require no genetic or optical access. This recording technique is also easily scalable to many cortical areas simultaneously (Steinmetz et al., 2019; Siegle & Jia et al. 2021; Chen & Liu et al. 2024). However, in these experiments, the cellular identities of the recorded neurons are almost entirely invisible. While atlases for identifying cell types from their transcriptomic expression are available (Tasic et al., 2018; Bakken et al., 2021), equivalent “cell type classifiers” that can match cell type to specific *in vivo* electrophysiological properties do not exist. This is because the correspondence between cell types and their *in vivo* electrophysiological properties are not understood.

Classically, one solution was to use one-dimensional measures such as the action potential width to differentiate “broad-spiking” putatively excitatory from “narrow-spiking” putatively fast-spiking inhibitory neurons (Mitchell et al., 2007; Hussar and Pasternak, 2009). However, such a division neglects differences in waveform shape important for differentiating cell types (Lee et al., 2021; Amatrudo et al., 2012; Povysheva et al., 2013; Krimer et al., 2005; Zaitsev et al., 2008) which is ultimately the result of transcriptomic differences in the expression of ion channels (Bomkamp et al., 2019; Bakken et al., 2021). Another notable electrophysiological property that differs between cell types is a neuron’s intrinsic firing rate pattern called its inter-spike interval (ISI) distribution (Latuske et al., 2015; Schneider et al., 2023). In addition, the circuit connectivity of various cell types is often distinct leading to differences in the input that a cell receives (Zaitsev et al., 2008). These differences in input, in combination with a neuron’s intrinsic dynamics, shape firing rate pattern in responses to external stimuli and task events also known as a peri-stimulus time histogram (PSTH) (Pinto and Dan, 2015; Yu et al., 2019; Ramadan et al., 2022). Thus, each modality provides different but complementary information about a cell and therefore an ideal approach would be to combine these different modalities according to their reliability or informativeness in delineating differences in cell type. However, it’s unclear which of these “electrophysiological modalities” are most important or else how they can be combined to create cell type classifiers that allow for the monitoring of multiple cell types *in vivo*. Therefore, optimally combining the information present across different modalities is critical for building electrophysiological cell type classifiers and ultimately, uncovering the function of neural circuits.

Here we present “PhysMAP”, an approach that reliably identifies different cell types by multi-modal integration of different physiological modalities such as a neuron’s waveform shape, ISI distribution, PSTH, derived electrophysiological metrics, etc. PhysMAP achieves multi-modal combination by combining non-linear dimensionality reduction (McInnes et al., 2018) with a weighted-nearest neighbor graph (WNN) construction from Seurat v4, a tool that has seen considerable success in multiomics (Hao & Hao et al. 2021). PhysMAP offers three key features: First, PhysMAP can incorporate any number of electrophysiological properties and arrive at a representation that captures underlying cell types better than any one modality (such as waveform shape) alone. Second, structure elicited by PhysMAP separates neurons by cell type and can delineate certain cell types in a manner that we demonstrate adheres closely to high-level “ground truth” labels available through optogenetic tagging (Yu et al., 2019; Lakunina et al., 2020; Petersen et al., 2021). Third, PhysMAP has the ability to learn a representation and infer cell types in new, unlabeled data allowing it to function as a cell type classifier. Thus, PhysMAP unlocks the study of circuit dynamics of multiple neuronal cell types simultaneously during sensation, perception, and action.

## Results

### Multi-modal analysis of extracellular electrophysiology with PhysMAP

Our PhysMAP approach uses a weighted graph combination solution developed in multiomics for intelligently integrating multiple modalities (transcriptomic, epigenomic, and proteomic data) in order to find cell type structure that transcends single modalities (Hao et al., 2021). As in our previous WaveMAP approach (Lee et al., 2021, 2023), PhysMAP uses the average normalized data for each single unit. Unlike WaveMAP, PhysMAP looks at arbitrarily many modalities beyond simply waveform shape. PhysMAP then finds shared underlying structure by combining these modalities via a weighted nearest neighbor (WNN) graph. After assessing all modalities for all cells, this method unifies all modality-specific graphs weighing each according to their informativeness on a per-unit basis. This graph is then projected into two dimensions for visualization of high-dimensional multi-modal structure. In Fig. 1, we show how this WNN graph is constructed in the simple two-modality case combining waveform shape and ISI distribution. What follows here is a description of these steps appended by the motivation for each. These steps are also described in greater detail in Methods: Weighted Nearest Neighbors.

**Figure 1.**
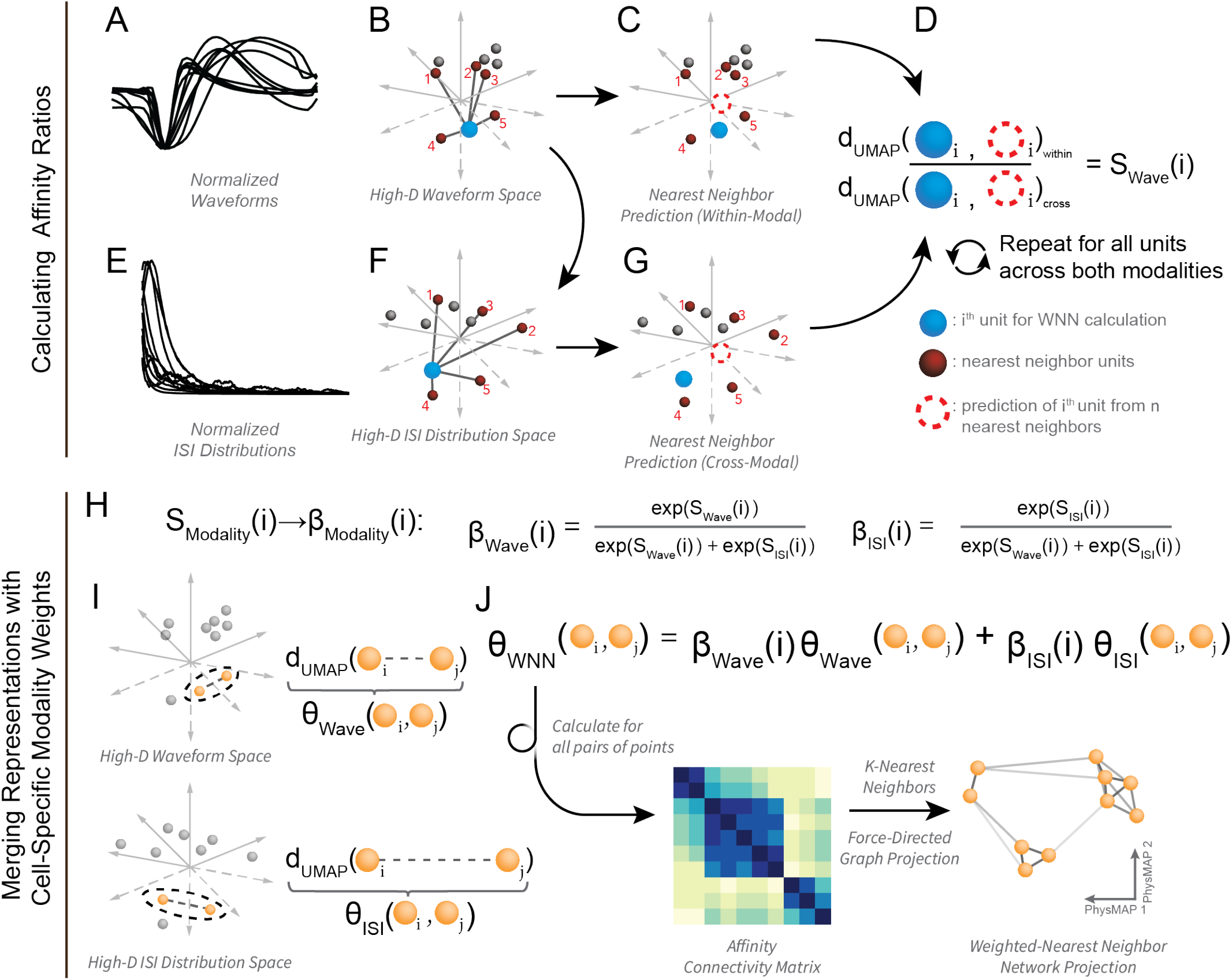
Schematic of PhysMAP and weighted-nearest neighbors algorithm. (**A**) A sample of normalized average single unit extracellular action potential waveforms. (**B**) Waveforms in (A) are shown in high-dimensional space with each axis pertaining to each time point along the waveform’s trace. For a sample unit (blue sphere), its nearest neighbor waveforms are highlighted (numbered red spheres). (**C**) The nearest neighbor waveforms are averaged to generate a prediction of the original waveform (dashed red circle). (**D**) The numerator of the waveform affinity ratio (S_Wave_(*i*)), the “within-modal affinity”, is taken as the difference between the original neuron’s average waveform and its nearest neighbor prediction. This “reconstruction error” distance is passed through a modified UMAP distance kernel. (**E**) Normalized ISI distributions drawn from the same neurons that produced the waveforms in (A). (**F**) The same neuron in (B; blue sphere) and its same nearest neighbors in waveform space are now shown in ISI distribution space. (**G**) As in (C), a prediction is made for the sample unit but this time in ISI-space (red dashed circle). The difference between these, passed again through a modified UMAP distance kernel, is used to form the “cross-modal affinity” that appears in the denominator of the affinity ratio in (D). The ratio of within- and cross-modal affinities form the waveform affinity ratio (S_Wave_(*i*)) for the sample unit. This ratio is calculated for each unit’s waveform and ISI affinity ratios (S_Wave_(*i*) and S_ISI_(*i*)). (**H**) Per-unit affinity ratios (S_Modality_(*i*)) for each modality are converted into per-unit modality weights (*β*_Wave_(*i*) and *β*_ISI_(*i*)) via these formulae. (**I, top**) The difference between a sample pair of points in high-dimensional waveform space is another type of “affinity” (*θ*_Wave_(*i, j*)). This is found by passing the distance into UMAP’s distance kernel as before. (**I, bottom**) The same pair of points as in (I, top) with the UMAP distance between them in high-dimensional ISI space (*θ*_ISI_(*i, j*)). (**J**) The equation for calculating weighted affinities (*θ*_WNN_) as a linear combination used in constructing the weighted-nearest neighbors graph. This weighted average of affinities is calculated for all points to construct an affinity connectivity matrix. The final multi-modal projection is found after applying *k*-nearest neighbors to the affinity connectivity matrix and projecting into two dimensions with UMAP’s projection algorithm (a force-directed graph layout procedure).

1. **Within-Modal Affinity**: Within the waveform space (Fig. 1A), we select a single neuron (blue sphere in Fig. 1B), identify its *k*-nearest neighbors (in this example, *k*=5; red spheres in Fig. 1B), and average them to predict the waveform of said neuron (dashed red circle in Fig. 1C). We then calculate the *within-modal affinity* by passing the neuron’s actual waveform and its predicted waveform into a modified UMAP distance kernel (Fig. 1D, numerator), and thus a measure of how well a neuron’s waveform is predicted by its neighbors in waveform-space. In the third step, we will show how this will be useful when determining which modalities to “trust” more in the form of an “affinity ratio”.
2. **Cross-Modal Affinity**: Next, we calculate a *cross-modal affinity* for the ISI distribution modality (Fig. 1E). Using the same neuron and its same neighbors in Fig. 1B, but now in ISI-space where each dimension is a time point along an ISI distribution curve (Fig. 1F), we calculate a predicted ISI distribution for this neuron by averaging (dashed red circle in Fig. 1G). We do this again by passing the neuron’s true ISI distribution and it’s predicted ISI distribution into a modified UMAP distance kernel (denominator of Fig. 1D), providing a measure of how well one modality versus another is predictive of a neuron’s properties using the same neighbors as to facilitate comparison.
3. **Affinity Ratio**: We define the per-unit *waveform affinity ratio*, S_Wave_(*i*), (fraction in Fig. 1D) as the ratio of this neuron’s waveform *within-modal affinity* divided by its ISI dist. *cross-modal affinity*. Conversely, the ISI distribution per-unit affinity ratio S_ISI_(*i*) is also obtained (not shown) by considering the ISI distribution-space as the within-modality and the waveform shape-space as the cross-modality. This ratio compares how well one modality versus another is predictive of a neuron’s properties and will be used to determine how much to weigh each modality for this neuron.
4. **Modality Weight:** We then calculate the weight associated with a unit for a particular modality (*β*_Modality_(*i*)) as the ratio of the exponentiated affinity ratio for a modality divided by the sum of exponentiated affinity ratios over all modalities (Fig. 1H; left equation for waveforms, right equation for ISI dists.). Thus for each neuron, each modality is differentially weighted according to how well the cell’s properties are predicted in each.
5. **Pair-Wise Unit Distances**: The distances between every pair of points (in every modality) is also passed into the same modified UMAP distance metric for construction of each modality-specific graph before re-weighting (Fig. 1I). For example, the edge between a pair of units *i* and *j* in waveform shape-space is *θ*_Wave_(*i, j*). In this way, subsequent calculations use UMAP’s notion of distance which has been shown to capture underlying low-dimensional manifold structure (McInnes et al., 2018).
6. **WNN Construction and Visualization**: Finally, we calculate a connectivity matrix of new pair-wise edges by taking the weighted linear combination of edges in the waveform space and ISI space with the pair-wise distances and affinity weights calculated in the previous steps (Fig. 1J). The final WNN graph is derived from this matrix by using *k*-nearest neighbors algorithm with *k* set to a large number (here it is 200). Using UMAP’s force-directed graph layout procedure, we then visualize the high-dimensional multi-modal structure by projecting the graph into two dimensions.

These steps proceed similarly in the case of three or more modalities with affinity ratios calculated for each modality against every other. The summations in the denominators of Fig. 1H and weighted average of Fig. 1J expand to include these other modalities as well. In the next sections, we use this PhysMAP approach and show how it can combine multiple modalities to delineate cell types in three different datasets.

### PhysMAP combines electrophysiological modalities to uncover cell type and laminar structure

*Ex vivo* experiments have shown that the relationship between cell type and electrophysiology spans many different physiological properties (Gouwens et al., 2020; Krimer et al., 2005; Zaitsev et al., 2008). These results are very promising and suggest that cell types could, in principle, be identified purely from modalities available from *in vivo* extracellular electrophysiological recording.

We examined if combining multiple electrophysiological modalities can find a structure that aligns to cell types better than any modality alone by applying PhysMAP to *in vivo* juxtacellular recordings from the mouse somatosensory (whisker barrel) cortex (Yu et al., 2019). Juxtacellular recordings provide low noise and well-isolated somatic waveforms recorded unambiguously from single neurons. This dataset also included anatomical and molecular discrete cell type information obtained through measured probe insertions, optogenetic tagging, and immunohistochemistry (Kvitsiani et al., 2013) allowing us to assess if PhysMAP can produce representations that align with ground-truth labels.

We first examined the two-dimensional embeddings produced by applying UMAP to these modalities individually (roughly equivalent to applying PhysMAP to single modalities). We found that UMAP applied to waveform shape (the approach taken by WaveMAP; Lee et al. 2021, 2023), led to embeddings that align quite well with underlying cell types and laminar structure (Fig. 2A) as each unit, labeled by cell type and layer, segregates accordingly. However, in this waveform modality, some cell types (such as layer 4 excitatory cells [E-4], in orange) are split in their structure. This split disappears in other modalities such as PSTH (Fig. 2B) or ISI distribution (Fig. 2C). In these other two representations, the structure delineating certain other cell types are lost. For example, PV^+^and SOM^+^ cells are less clearly delineated from other populations in both the PSTH and ISI distribution visualizations. These results suggest that different modalities provide differing views into underlying structure and perhaps, by optimally combining several modalities, one can find a more robust delineation of cell types.

**Figure 2.**
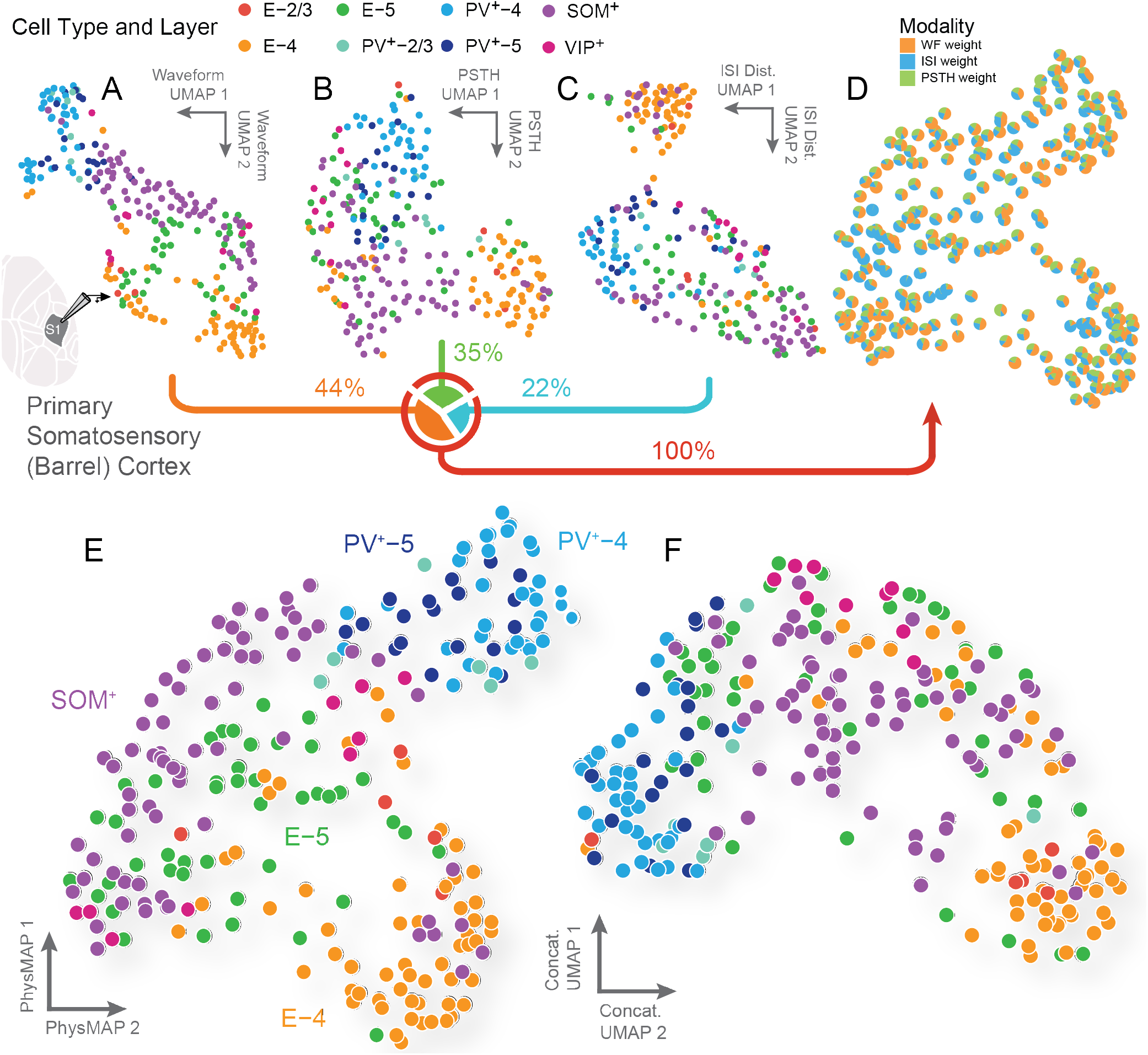
Multi-modal combination of physiological properties leads to structured representations that align with cell types. (**A**) UMAP on normalized average waveform shapes of neurons recorded *in vivo* juxtacellularly from mouse primary somatosensory cortex (Yu et al., 2019). Each unit is colored according to its ground truth cell type and layer listed above. Each modality was combined in different proportion according to the sum of each modality’s per-unit contribution to the total; this total contribution is listed as a percentage below each scatter plot. (**B**) UMAP on ISI distributions of the same neurons in (A). (**C**) UMAP on the PSTH firing rate distributions from each neuron in response to a deflection of a whisker during whisking. (**D**) Each neuron in PhysMAP’s WNN representation weighs each modality differently; these modal contributions are the sum of nearest neighbor edge modality weights (*β*_Modality_ in Fig. 1J) and are shown as proportions of each pie chart. (**E**) Each of the three modalities are integrated into one representation using the weighted nearest neighbors approach (Hao & Hao et al. 2021) and shown. Again, each neuron is colored according to their ground truth cell type and layer. (**F**) All three modalities in the dataset from Yu et al. (2019) were concatenated into a single data vector per unit and passed into UMAP. This represents each modality in unweighted combination (as opposed to the WNN approach in PhysMAP). Each neuron in the projected representation is again labeled according to cell type and layer.

To assess if multimodal combination is useful, we examined the two-dimensional representation of PhysMAP colored according to cell type and layer. We find that a combined representation obtained by weighting each modality on a per-neuron basis appears to best separate cell types (Fig. 2E). The resultant multi-modal representation also offers additional insight even in the two-dimensional projection: In this visualization, we see that fast-spiking parvalbumin-positive cells (PV^+^) are tightly clustered and segregated by layer. This visualization also shows that while many SOM^+^ cells are separable they occupy a large area in the PhysMAP representation which implies underlying physiological diversity (Urban-Ciecko and Barth, 2016; Wu et al., 2023; Yavorska and Wehr, 2016). Similarly, we observe that excitatory cells are highly distinct between layers 4 and 5.

To gain insight into how the WNN approach works, we next examined how PhysMAP combines each of these modalities on a per-neuron basis. Fig. 2D plots the proportion of each modality used for each data point in Fig. 2A, B, and C after combining all three modalities in PhysMAP and projecting into two dimensions. For the majority of units, waveform shape is the most informative modality as shown by the predominance of its color (orange) across most units. Tallying the contribution of waveform shape, it composes 44% of the total as opposed to PSTH which provides 35% and ISI dist. with 22%. However, there are certain regions of the projection in which other modalities (such as ISI distribution at bottom center of the plot) are more important. Together, these visualizations and results suggest, but do not confirm, that PhysMAP takes a nuanced per-neuron approach to combining modalities and this may help better separate cell types.

One concern is that perhaps our PhysMAP representations are a trivial result of having more information through the simultaneous inclusion of multiple modalities. To test if this was the case, we concatenated all three modalities into a single feature vector and ran PhysMAP on this concatenated “super property” which can also be thought of as all three modalities in unweighted combination (Fig. 2F). We found that the representation obtained by simple concatenation of the physiological properties alone demonstrated far weaker structure than the one found in PhysMAP (compare Fig. 2F to Fig. 2E). E-5 and SOM^+^ neurons are more intermixed with one another and also with PV^+^ types. Furthermore, all cell types form less well-structured clusters. This visualization shows that PhysMAP’s improvement in segregating neurons by cell types and layer is not a trivial result due to more information and instead that per-neuron weighted averaging is better in making use of multiple modalities.

We emphasize that the two-dimensional representations in this section serve purely as *visualizations* of high-dimensional structure and as such we only examine them here qualitatively. In the following sections, we analyze higher-dimensional representations directly to perform quantitative comparisons.

### PhysMAP delineates cell types better than any modality alone or in unweighted combination

The results from the previous section showed that, visually at least, the multi-modal representations obtained from PhysMAP align better with cell types than representations generated by individual modalities alone or in unweighted combination. To more precisely quantify these results in each case (uni-modal, unweighted multi-modal, and weighted multi-modal), we conducted two analyses, one supervised with a classifier and one unsupervised with clustering.

In a supervised analysis, we trained a classifier to identify each cell type using either the individual modality representations, the unweighted concatenation, or the weighted PhysMAP representation. For comparison, we also trained classifiers on derived electrophysiological metrics and the full dimensional raw data without dimensionality reduction. See Methods: Classifier Analysis for more details. For these classification analyses, we projected the WNN or unimodal UMAP graphs into a 10-dimensional embedding space for each of the various modality representations. We conducted an 15-85% test-train split and trained a gradient boosted tree model (GBM) with five-fold cross-validation on this dataset to identify one of five cell types that had over 25 examples each and this process was repeated 10 times with different random seeds. For each of the weighted multi-modal (PhysMAP), unweighted multi-modal (concatenation), and three individual uni-modal representations, we trained this classifier on a 10-dimensional embedding of the WNN graph. We plotted the performance of each of these classifiers trained on the high-dimensional representations against each other. PhysMAP matched or exceeded the performance of all other modalities singularly or in unweighted combination for all cell types (Fig. 3A), fully consistent with the visualizations in Fig. 2D and E,. We point out that the waveform modality (WF) performed nearly as well as PhysMAP and equating performance for three cell types (PV^+^-4, E-4, and SOM^+^) which confirms our previous observation that waveform shape is the most cell type informative modality (Fig. 2D). This finding of waveform shape’s importance in classifying cell types also serves as validation of our previous approach and findings with WaveMAP (Lee et al., 2021). The *benefit of adding the additional modalities comes from the improved classifiability of the E-5 and PV*^+^*-5 neurons from the rest, which was much poorer with only waveforms*.

**Figure 3.**
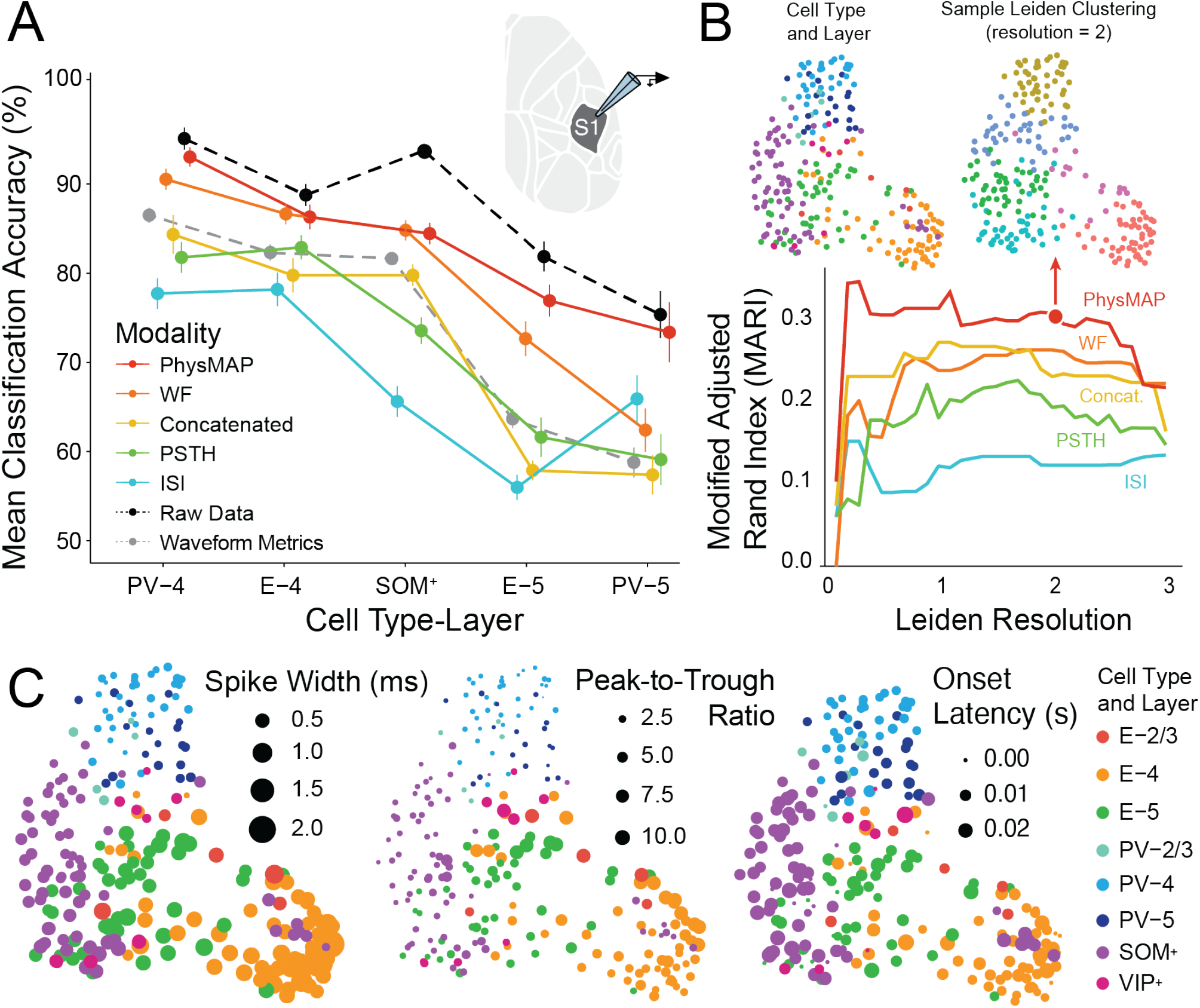
PhysMAP identifies cell types from juxtacellular recordings better than any modality alone. (**A**) A gradient boosted tree model (GBM) classifier was trained with 5-fold cross-validation on the 10-dimensional projection of each modality’s UMAP graph individually or on the 10-dim. projection of the multi-modal WNN graph. The same classifier was also trained directly on the full data (that is, without constructing a UMAP graph and projecting it), on the 10-dim. projection of the UMAP graph of the data concatenated into a single feature, and on the two derived waveform metrics (spike width and peak-to-trough ratio). The balanced accuracy performance of this classifier (mean *±* S.E.M.) on held-out data for each modality, combined modalities, and derived metrics is shown for the five cell type classes with over 25 units each. (**B, top**) The PhysMAP projection of neurons with their ground truth cell type and layer (left) next to an example Leiden clustering (Traag et al., 2019) with resolution parameter set to 2. (**B, bottom**) Leiden clustering is applied to the UMAP graphs of each modality alone (waveform [WF], ISI dist., and PSTH), in weighted (PhysMAP), or unweighted combination (concat.) and shown with the associated modified adjusted Rand index (MARI; Sundqvist et al. (2020)) calculated across a range of resolution parameter values from 0.1 to 3.0 in 0.1 step increments. Arrow marker indicates the clustering on PhysMAP used above in B. Waveform metrics and raw data were omitted in this analysis because these do not yield a high-dimensional UMAP graph. (**C**) The spike width, peak-to-trough ratio, and spiking onset latency for each neuron are shown via log transformed marker size under PhysMAP’s two-dimensional projection.

PhysMAP also demonstrated better classification performance than the modalities in an unweighted combination (concatenated) for all cell types. Similar to visualizations comparing the embeddings of the weighted and un-weighted modalities in combination (Fig. 2E vs Fig. 2F), our classification analysis demonstrates that PhysMAP is able to draw out cell type structure in a manner that is not trivially the result of including several modalities simultaneously.

Finally, PhysMAP (and the full waveform) far outperform derived waveform metrics which are how cell types are traditionally identified. This is a result that is consistent with general notions in machine learning that hand-derived metrics, while seemingly “interpretable” can lose important information. In our case features do not fully capture the full diversity of waveform shape in a way that is useful for identifying cell types.

#### Control

Classification performance from PhysMAP was robust to both the embedding dimensionality chosen for the WNN graph. First, based on a linear estimate of intrinsic dimensionality using principal component analysis, a 10-dimensional embedding was chosen for the classification analysis given that the “elbow”, normally regarded as the sufficient number of dimensions to capture dataset variance, was at the third, fourth, and third PC’s (which sum to 10) representing 91%, 80%, and 93% of the total variance for waveform shape, ISI dist., and PSTH respectively (Fig. S1). However, we also found that the particular choice of embedding dimensionality actually did not have a large effect on classifier performance. We examined the full range of embedding dimensionalities from 30 (matching the lowest ambient dimensionality of the input modalities) down to 2 (the plot Fig. 2E itself) and found very similar patterns of classification accuracy across all cell types and all modalities (Fig. S2A).

#### Control

Similarly, the particular classification algorithm used had little effect on this classification analysis. We also performed this supervised classification analysis across five other classifiers: a random forest, radial basis support vector machine (rbSVM), a classification tree, a *k*-neighbor nearest classifier, and a neural network with single hidden layer. Each showed very similar results demonstrating that our results are not classifier-dependent (Fig. S2B). Thus, in Fig. 3A we demonstrate that PhysMAP is able to intelligently combine multiple modalities in a manner that draws better cell type delineations than any modality alone or in unweighted combination.

#### Comparison

One potential criticism is that the interpretability of our PhysMAP approach comes with a cost and that some information is lost. However, PhysMAP performs nearly as well as a classifier on the raw data in detecting cell types. When the GBM classifier is trained directly on the full dimensional dataset from all modalities directly (that is, not on a low-dim. projection of the UMAP graph), performance is not much better on most cell types except for SOM^+^ cells (Fig. 2A, dashed line). Furthermore, training a classifier directly on the data loses all interpretability since it only returns a probability that some neuron belongs to a given class. Thus, while there is some information loss because of PhysMAP’s dimensionality reduction, it still produces a representation that contains most of the information necessary for both cell type classification. In return it provides a useful visualization for exploring physiological diversity. We will explore how this is useful in the next section.

### PhysMAP is also better in unsupervised settings

We also examined if PhysMAP representations aligns to underlying cell types even when no labels are available to train on. In most electrophysiological datasets, only a subset of the recorded cells will have ground truth labels. In a fully-unsupervised analysis, we asked the question of “without access to *any* ground truth, how well do unsupervised methods identify cell types?”. Thus, by examining the representations produced by PhysMAP in the extrema of having both complete ground truth or zero ground truth (supervised vs. unsupervised algorithms), we gain a more complete picture of PhysMAP’s performance over a range of possible dataset conditions.

We assessed how well unsupervised clusters aligned to underlying ground truth cell types by calculating the modified adjusted Rand index (MARI; Sundqvist et al. (2020)) between each. This is a measure of how often the same ground truth cell types co-cluster under an unsupervised clustering. We used cluster labels generated from Leiden clustering applied to the high-dimensional graphs produced by PhysMAP and each physiological modality (that is, the graphs before projection) across a range of resolutions (cluster sizes) and calculated the MARI score between the two sets of labels. See Methods: MARI Calculation for more details. The greater the MARI value, the more the clusters contain singular cell types each. To provide an example of how Leiden clustering on PhysMAP compares to ground truth, we show neurons in PhysMAP colored according to cell type (Fig. 3B, top left) alongside a sample Leiden clustering with resolution set to 2 (Fig. 3B, top right). Examining each resolution parameter of Leiden clustering, PhysMAP produced higher MARI scores than any electrophysiological modality alone or in unweighted combination for all Leiden resolutions (Fig. 3B). This performance was followed by modalities in unweighted combination and waveform shape alone, PSTH, and then ISI distribution. Collectively, these results demonstrate that PhysMAP is useful in identifying cell types in both supervised and unsupervised settings.

### PhysMAP provides interpretable representations and biological insight

Thus far, we have demonstrated that PhysMAP outperforms individual modalities at uncovering the link between electrophysiological properties and underlying cell types in supervised and unsupervised settings. However, it is increasingly clear that cell types are not “discrete” and physiological variation within a cell class is the norm and not the exception (Cembrowski and Menon, 2018). Ideally, any tool for cell type identification should allow one to visualize this diversity, both within and across discrete cell types.

We found that PhysMAP’s visualizations were ideal for this purpose of examining underlying diversity. Fig. 3C, shows the PhysMAP plot in Fig. 2E but now with marker radius scaled to either reflect waveform spike width, peak-to-trough ratio, and onset latency. Examining spike width Fig. 3C, left, many trends immediately pop out: PV^+^ cells seem to be uniformly narrow no matter the layer; SOM^+^ cells occupy a gradient of widths and overlap with E-5 cells at their widest; and the widest spiking cells are E-4. This finding regarding SOM^+^ cells occupying a range of spike widths is likely because it is composed of several subtypes of varying widths (Hostetler et al., 2023) which have distinct computational roles (Wu et al., 2023). Waveform peak-to-trough ratios also differ between cell types Fig. 3C, middle with PV^+^ and SOM^+^ cells having the smallest and both E-4 and E-5 cells occupying a range which includes the largest ratios. Finally, we replicated that onset latency (half the time it takes to reach peak firing rate after whisker deflection; Fig. 3C, right) is fastest for PV^+^ cells and SOM^+^ cells are some of the slowest, consistent with the reports from the original study (Yu et al., 2019).

These insights were less clear than when only examining the metrics without using the PhysMAP representation. We examined a scatter plot of the spike width and peak-to-trough-ratio to assess whether any structure easily pops out (Fig. S3). While the scatter plot shows some cell type structure it appears like one continuous relationship and fails to show any easy separability between SOM^+^, PV^+^ and excitatory cells. In addition, in either waveform shape metric, the distributions of the the PV^+^ and excitatory cells of different layers overlap as shown by the coinciding PV^+^-4/PV^+^-5 or E-4/E-5 histograms on the marginals of Fig. S3.

In summary, PhysMAP provides the ability to easily and simply combine multimodal electrophysiological data to obtain a representation that 1) classifies cell types better than any individual modality or feature alone, 2) can also be used in unsupervised settings to separate out candidate cell types, and 3) provides an interpretable visualization that can be used for additional exploration and development of hypotheses about data.

### PhysMAP uncovers cell types in extracellular electrophysiological recordings

The juxtacellular recordings analyzed in the previous section represent a best case scenario for PhysMAP for several reasons. First, this recording method offers access to somatic waveforms that are unperturbed by distance to spike origination site or from differences in spike shape occurring due to the differential ion channel properties of neurites (axons and dendrites) (Someck et al., 2023). Second, juxtacellular recordings offer unambiguous isolation of single unit activity. In contrast, extracellular recordings are prone to poor isolation which blurs the properties of different cell types (Vincent and Economo, 2023). This results in many multi-unit “cell types” occupying intermediary properties. Third, juxtacellular recordings allow for the preferential targeting of particular cell types that may be less prevalent. Inhibitory types make up only 20% of neurons in mouse cortex (Tremblay et al., 2016) but in this set of juxtacellular recordings they compose 56% of the dataset as they have been preferentially targeted for recordings. This provides the downstream benefit in that a classifier will not be biased towards more prevalent cell types or will fail to classify less common ones. We asked whether PhysMAP could identify find cell types even when applied to recordings acquired via extracellular probes. To address this question, we first analyzed an extracellular electrophysiological dataset collected from mouse primary auditory cortex (A1, Lakunina et al., 2020) with optotagged cells.

In this experiment, Lakunina et al. (2020) recorded from neurons in A1 with extracellular probes while mice were presented with auditory stimuli. They identified both SOM^+^ and PV^+^ units using optogenetic tagging and recorded each unit’s waveform shape and ISI distribution (over 50 ms windows). We passed these two modalities into PhysMAP to produce a combined representation (Fig. 4A). In this projected space, both SOM^+^ and PV^+^ cell types (purple and teal respectively) are well-separated and occupy a distinct region of the embedding that clusters away from a much larger number of “untagged” (presumably excitatory) cells (in gray, Fig. 4A). This result complements the findings in the previous juxtacellular dataset (Yu et al., 2019) and confirms that PhysMAP is able to separate out cell types even in an extracellular setting.

**Figure 4.**
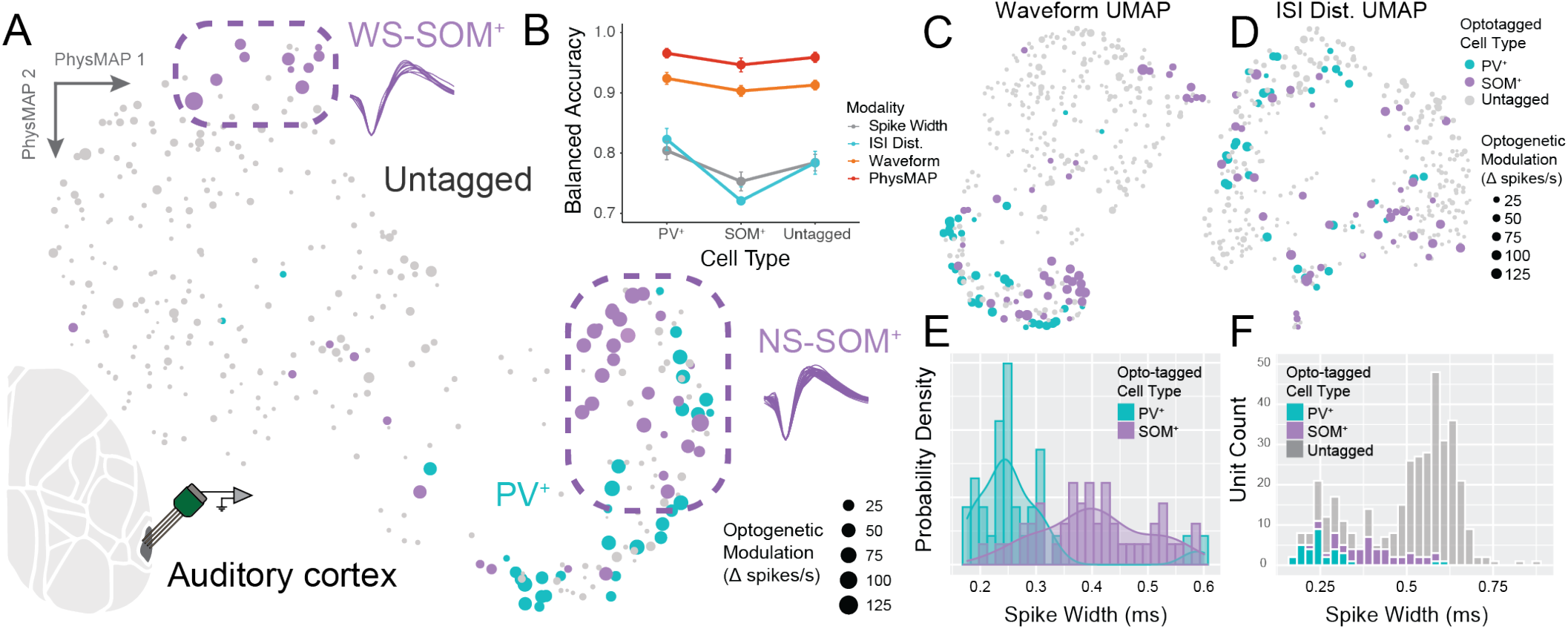
PhysMAP identifies cell types in extracellular recordings. (**A**) PhysMAP applied to extracellular recordings from mouse auditory cortex (Lakunina et al., 2020) using silicon probes. We used waveform shape and ISI distribution as our modalities and neurons are colored according to cell type whose identities were obtained via optogenetic tagging. Marker size is set according to the increase in spike rate from optogenetic stimulation. Highlighted are a population of wide-spiking SOM^+^ cells (top) and narrow-spiking SOM^+^ cells (right). (**B**) A GBM classifier with five-fold cross-validation was trained on the 10-dimensional graph of each modality, PhysMAP, or waveform spike width to identify each optotagged cell type or untagged label. The balanced classification accuracy is shown (mean *±* S.E.M.) with many error bars smaller than marker size. (**C**) Normalized average waveforms in this dataset are passed into UMAP and their projection shown with optotagged cells colored and degree of optogenetic modulation (average change in spikes/s during stimulation epochs) shown by marker size. (**D**) Similarly, ISI distributions for these same neurons are also passed into UMAP and their projection shown. (**E**) Probability density estimates for the distribution of each optotagged type across a range of spike widths is shown both as a histogram and with a kernel density estimator (solid line). (**F**) Additionally, spike width is shown for all cells, including untagged ones.

We again examined if PhysMAP better separates out these cell types than any single modality or waveform metrics alone. As in the previous section, we quantified this separation by training a classifier to identify each cell type using PhysMAP and benchmarked this classifier’s performance against classifiers trained on individual modalities (waveform shape or ISI distribution) or waveform metrics (spike width). Note that MARI was not calculated for this dataset as such an analysis is only valid when all labels are known. The classifier on PhysMAP identified PV^+^ and SOM^+^ cells with high accuracy (97% and 95% mean balanced accuracy, respectively) and surpassed classification performance using ISI distributions or waveform metrics (82% and 72% or 80% and 75% mean balanced accuracy, respectively), the latter of which are how cell types are identified in much of the literature (Fig. 4B). PhysMAP also slightly outperformed classification on waveform shape for both PV^+^ and SOM^+^ cells (92% and 90% mean balanced accuracy, respectively). The reason for this near-equivalence is that waveform shape is highly informative of underlying cell types and thus PhysMAP correctly prioritizes this modality. The median unit weight associated with the waveform shape and ISI distribution modalities were 0.71 and 0.29 respectively. This prioritization of waveform shape over ISI distribution was also evident in their respective UMAP visualizations: the PhysMAP’s projection appears almost exactly the same as UMAP on waveform shape (Fig. 4C). In contrast, a UMAP applied to ISI distribution shows far less structure Fig. 4D. Furthermore, although these does not correspond to a particular cell type, classification with PhysMAP outperforms any other modality in identifying “untagged” cells. By mirroring results from our previous analysis of juxtacellular recordings, we establish that PhysMAP is also capable of uncovering cell types in extracellular recording settings.

Again, as in the juxtacellular dataset, the PhysMAP approach provided interpretable representations. To illustrate this point, we identified two sub-populations of SOM^+^ cells not documented in the original publication of this dataset (Lakunina et al., 2020) but predicted by the literature (Hostetler et al., 2023; Wu et al., 2023; Urban-Ciecko and Barth, 2016). These wide-spiking SOM^+^ (WS-SOM^+^, Fig. 4A, upper) and narrow-spiking SOM^+^ (NS-SOM^+^, Fig. 4A, lower right) sub-populations occupied separate regions of the PhysMAP embedding and differed significantly in their spike width (0.48 *±* 0.04 vs. 0.37 *±* 0.05 ms [median *±* S.D.], respectively; p *<* 0.001, Mann-Whitney U test). These sub-populations have been observed in other experiments to pertain to SOM^+^ cellular subtypes with specific synaptic targeting (Wu et al., 2023) and differential function during behavior (Kvitsiani et al., 2013; Kim et al., 2016).

Note that these two sub-populations are again not trivially identifiable from spike width alone: SOM^+^ cells, as a whole, were part of a continuum under this metric (Fig. 4E, in purple) and overlapped with PV^+^ cells (Fig. 4E, in teal) in width. Even if an arbitrary width cutoff was used to bisect SOM^+^ cells into two sub-populations, the WS-SOM^+^ cells would occupy a range of widths that strongly coincides with large number of other untagged— presumably excitatory—cell types (Fig. 4F) and thus would not be able to be identified uniquely. Only PhysMAP is able to simultaneously disambiguate these two SOM^+^ subtypes in addition to PV^+^ and untagged putatively excitatory cells.

### PhysMAP finds the same cell types across heterogeneous datasets from different labs

We have demonstrated that PhysMAP uncovers cell types in datasets from single labs but it would be beneficial if a classifier could also find cell type structure across datasets gathered by different labs. Cell type classifiers could then be built by small collaborations of interested investigators across different labs. To this end, we combined extracellular recording datasets originating from separate labs each containing cells from both visual cortex and hippocampus (Siegle et al., 2021; Petersen et al., 2020) released as part of CellExplorer (Petersen et al., 2021).

For neurons from both datasets, we had measurements of waveform shape, ISI distribution, spike autocorrelogram (ACG), and eleven derived electrophysiological metrics such as spike width, coefficient of variation, and ACG delay time constant etc (full list in Methods: Extracellular Mouse Visual Cortex and Hippocampus Dataset). Similar to our two previous analyses, all modalities ostensibly showed cell type-dependent structure but differed in which cell types were better separated: waveform shape (Fig. 5A) splits PV^+^ cells into two regions whereas the other modalities do not. Similarly, pyramidal cells are well-isolated under ISI distribution (Fig. 5B) but less well in others like ACG (Fig. 5C). Derived electrophysiological metrics (Fig. 5D) present an intuitive PV^+^ to SOM^+^ to pyramidal cell structure but separate out SOM^+^ cells less than using the ISI distribution. PhysMAP combines these modalities to produce a representation that produces a clear separation of cell types (Fig. 5E).

**Figure 5.**
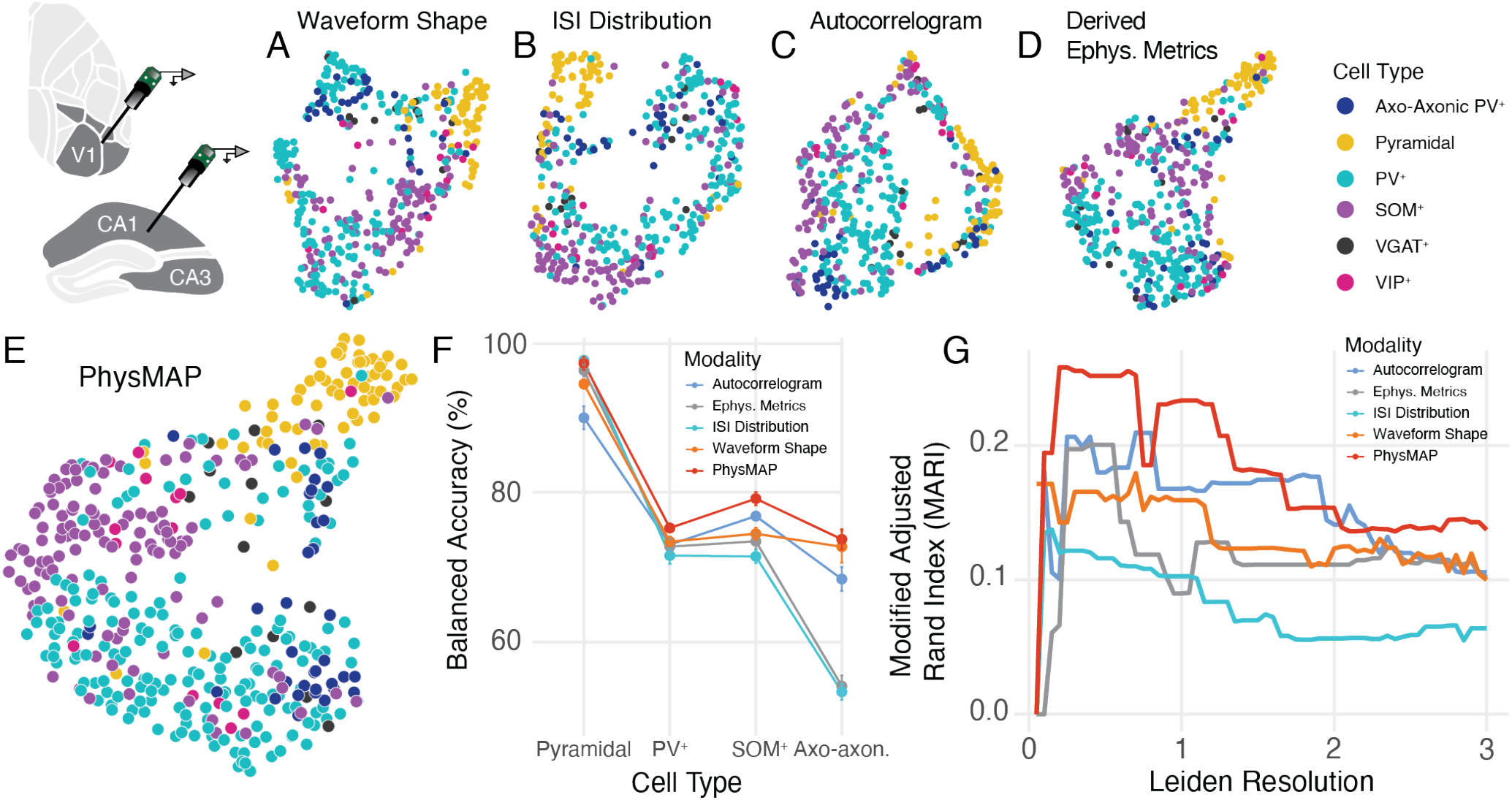
PhysMAP is able to identify cell types across areas and labs. (**A - D**) Across a pooled dataset of visual cortex and hippocampal extracellular recordings from CellExplorer, we show 2-D UMAP visualizations applied to waveform shape, inter-spike interval distribution, autocorrelogram, and thirteen various derived electrophysiological metrics respectively. Cell type classifications were obtained by opto-genetic tagging. (**E**) The previous four modalities are combined into a unified PhysMAP representation and visualized in 2D. (**F**) A gradient boosted tree model (GBM) classifier with 5-fold cross-validation was trained on the 10-dimensional projection of each modality’s high-dimensional UMAP graph or on PhysMAP’s multi-modal WNN graph. The balanced accuracy for the classifier is shown for each modality and cell type that contained more than 25 examples. (**G**) Leiden clustering is applied to each modality alone and to PhysMAP and the corresponding modified adjusted Rand indices are calculated across a range of resolution parameter values from 0.1 to 3.0 in 0.1 step increments. All cell type labels were included in this calculation.

In order to evaluate PhysMAP quantitatively for this dataset, we again calculated both the accuracy of a classifier and each representation’s MARI score. We again trained a GBM classifier with 5-fold cross-validation to identify the four cell types with more than 25 units each from the 10-dimensional embedding of each modality and PhysMAP. For all cell types (pyramidal, PV^+^, SOM^+^, and axo-axonic [a PV^+^ subtype]), PhysMAP provided the best classification accuracy (Fig. 5F).

Next we evaluated how well cell types clustered by calculating the MARI score of each representation. As before, across nearly all Leiden resolution values, PhysMAP provided the best unsupervised clustering of cell types (Fig. 5G). We also examined if hippocampal and cortical inhibitory cells were differentiable (Fig. S4A). Both PV^+^ (Fig. S4B) and SOM^+^ (Fig. S4C) cells did not form separate populations in PhysMAP regardless of the area despite being well-differentiated between types. Thus, we expect that that a cell type classifier will generalize across areas to some extent even between areas as different as hippocampus and neocortex. Thus, even when combining data from multiple labs and areas, PhysMAP better separates cell types than any single modality suggesting the feasibility of building cell type classifiers in a collaborative manner. We next evaluate how well PhysMAP performs as a cell type classifier.

### PhysMAP allows for the creation of a cell type classifier to accurately identify several major cell types

So far, we have shown that PhysMAP is better able to unsupervisedly identify cell types in three areas of mouse cortex and hippocampus but in these analyses, we compared modalities by training a classifier after data transformation to facilitate comparison with limited data sizes. In order to properly test PhysMAP’s functioning as a cell type classifier, we evaluate classification on held-out, un-transformed data. To build the cell type classifier, we took advantage of Seurat’s multi-modal reference mapping tool which allows for classification of cell types given some reference dataset. Here, we take the previous CellExplorer dataset and test PhysMAP’s ability to identify the identity of a random subset of the total dataset with the remainder used to build a “reference map”. With this analysis, we show that PhysMAP maintains good generalization on three of four major cell types available (excitatory pyramidal, PV^+^ interneurons, and SOM^+^ interneurons).

To begin, we consolidated the CellExplorer dataset such that axo-axonic PV^+^ chandelier cells were relabeled as “PV^+^” (Dudok et al., 2021) and removed VGAT^+^ neurons as they are pan-inhibitory and thus do not label one particular inhibitory cell type. We then divided the 417 resulting cells into an 15-85% test-train split. The training data—containing waveform, ISI distribution, and single derived metrics measuring the former two—was passed into PhysMAP to generate a reference map (Fig. 6A). A query dataset of held-out neurons was then mapped onto this reference using anchor point alignment (Haghverdi et al., 2018) and cross-projection of query data onto the reference map (Stuart & Butler et al. 2019). See Methods: Reference Mapping with Anchors and Cell Type Classification for methodological details.

**Figure 6.**
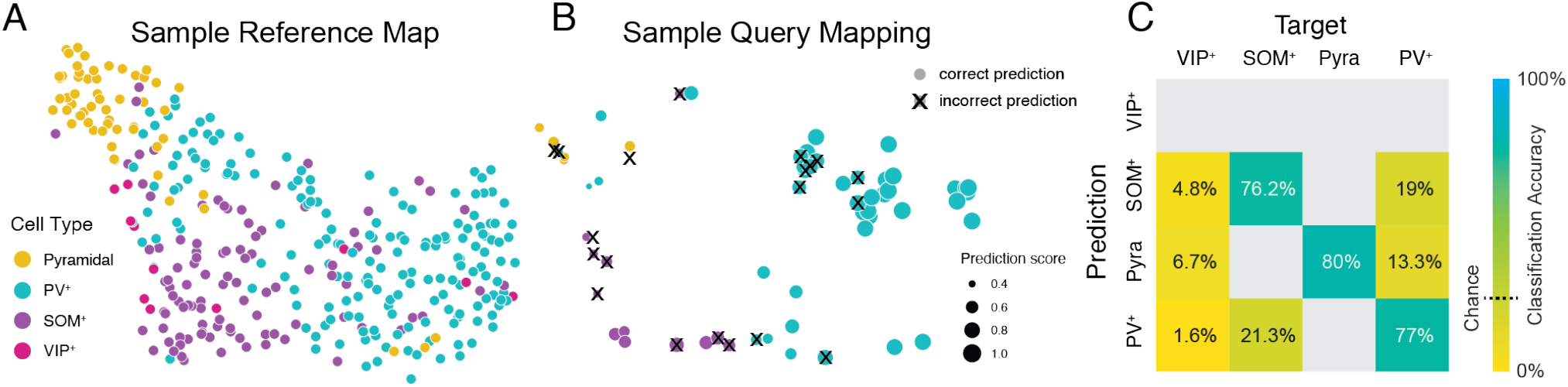
PhysMAP makes it possible to construct a cell type classifier for three major cell types. (**A**) Using the previous CellExplorer dataset, axo-axonic cells were relabeled as PV^+^ (because they are a PV^+^-subtype) and VGAT^+^ cells were omitted since they label inhibitory neurons as a whole. The four modalities in (Fig. 5A-D) were then passed into PhysMAP for a given training set and a 2-D visualization for this “reference map” is shown. (**B**) The held-out test set is then projected into the reference map space using anchor alignment and visualized as a “query mapping”; cell type labels are according to nearest neighbor predictions after projection. Incorrect predictions identified by a small “x” plotted over each data point. Each cell is also sized according to “prediction score” which is an empirical measure of the quality of nearby anchors; greater prediction scores imply a closer match between local query mapping and the reference map at that location. (**C**) A confusion matrix of query mapping predictions used as a classifier across the four major cell types in (A). Each row contains the mean percentage (after 10 instances of classifier training) that each predicted cell type label was assigned to its true target cell type. The percentage of cells that were correctly predicted is shown along the main diagonal. Some tiles are left blank because no predictions were made for that pair of predicted and target cell type.

Fig. 6B shows a sample query mapping of the test set colored according to predicted cell type with incorrect predictions overlaid with a black “x”. Cell type predictions are made based upon a query cell’s nearest neighbors in the reference map. These data points are also sized according to their “prediction score” which is a per-neuron measure of confidence in the anchor alignment; higher scores are associated with higher quality alignment. In this sample query mapping, neurons with low prediction score more often result in incorrect predictions. Thus, if a more conservative prediction of cell types is desired, only neurons with higher prediction scores can be used.

To evaluate the anticipated performance on this dataset with PhysMAP performing as a cell type classifier, we conducted ten random test-train splits of the dataset and examined the average classification performance. We found that although VIP^+^ cells were not identifiable (the query mapping never predicted them due to low sample number) all other cell types were highly identifiable (Fig. 6C). Each of SOM^+^, PV^+^, and excitatory pyramidal cells were identified at or above 75% accuracy which was far above chance (25%). In summary, we find PhysMAP not only locates more cell type structure than other modalities, it also provides a cell type classifier that can be used to identify multiple cell types from *in vivo* extracellular recordings.

## Discussion

Our goal in this study was to assess if a method to compute a weighted combination of electrophysiological modalities could identify underlying neuronal cell types and serve as a classifier. Our approach, PhysMAP, integrates electrophysiological modalities to produce a combined representation that separates out cell types better than any single modality alone or in unweighted combination. Specifically, in both juxtacellular and extracellular *in vivo* experiments, the multi-modal representation produced by PhysMAP was better at uncovering cell types in both unsupervised and supervised settings. We also demonstrated excellent classification performance in identifying three major cell types from held-out data in an interpretable manner.

We note that PhysMAP is not the only approach for merging information from multiple modalities. Other prominent tools in multiomics including MOFA (Argelaguet et al., 2020) and MOJITOO (Cheng et al., 2022) can perform similar integration of multiple modalities. We used the WNN algorithm from Seurat v4 to develop our PhysMAP approach because it has demonstrated excellent performance for multiomics problems (Cheng et al., 2022) and we found that this performance extends to identifying cell types when combining multiple electrophysiological modalities. We also used Seurat because it is perhaps the most extensively used package in the multiomics community, has a large community of contributors, and is continually being improved. For instance, Seurat v5 adds “bridge integration” allowing the prediction of a cell’s proteomic profile given its transcriptomic expression by means of a “dictionary” (Hao et al., 2023). Applied to an electrophysiological recording, one might envision predicting a neuron’s waveform shape given only knowledge of its ISI distribution.

We used the waveform, inter-spike interval, and peri-stimulus time histogram in PhysMAP to differentiate cell types. Other modalities might also be important for delineating cell types. For example, the spike-local field potential (spike-LFP) coupling has been posited to separate PV^+^ and SOM^+^ cell types (Wei et al., 2023; Onorato et al., 2023). In addition, there may be particular “localizer” stimuli that can be used to generate PSTHs to discriminate certain cell types. For example, a full-field flash has long been used to identify thalamocortical input cells into V1 (Heynen and Bear, 2001). However, one must be judicious about the modalities chosen because of circularity. That is, if a certain cellular property wants to be compared between cell types, it should not be passed into PhysMAP because such properties will already be used to select the groups to be compared (Kriegeskorte et al., 2009).

PhysMAP offers two advantages over optical imaging approaches: simultaneous monitoring of neuronal cell types on single trials and the exploration of cell types in primates. First, they allow for the *simultaneous* monitoring of neural populations, which is currently difficult in optical imaging experiments due to constraints on optics that limit imaging to only two fluorophores (Aharoni and Hoogland, 2019). Simultaneous monitoring will help better understand circuit dynamics on single trials (Peixoto et al., 2021; Boucher et al., 2023), and bridge the gap between anatomical microcircuits and dynamics (Esparza et al., 2023). Second, cell type-specific imaging often requires transgene delivery and expression. In contrast, PhysMAP only requires recordings of the electrical activity of single neurons and is therefore particularly useful for the study of cell type-specific neural dynamics in non-human primates (NHPs). NHPs are a valuable and scarce resource and each fully-trained animal can perform multiple sophisticated tasks. Primate researchers are therefore cautious in using direct cell type-specific recording techniques that entail risky procedures such as viral injections or cranial window installations. In addition, in primates, optical methods and genetic engineering tools are difficult to execute without enormous resources and the geometric constraints are even more complicated because of the size of cortex (Trautmann, O’Shea, & Sun et al. 2021, O’Shea et al., 2017). For this reason, PhysMAP would be a no risk solution to obtaining cell types when applied to routine electrophysiology experiments.

Naively, one might be tempted to question why dimensionality reduction methods like PhysMAP should be used at all. Instead, would a purely classification-based approach not suffice to identify cell types? While current classifiers will delineate some cell types from electrophysiological properties (Schneider & Azabou et al. 2023, Ye & Shelton et al. 2023), we find that these do not outperform PhysMAP in identifying the same cell types. More importantly, PhysMAP has two key advantages over **pure discrete classification** approaches.

First, PhysMAP produces interpretable visualizations that can reveal heterogeneity within and across cell types. This heterogeneity manifests as an overlap in the embeddings of one cell type over another in the PhysMAP visualization. For example, we observe considerable variability within SOM^+^ cell types in both juxtacellular and *in vivo* extracellular settings and across cortical areas. In both the mouse S1 and A1 datasets, we found that SOM^+^ cells were heterogeneous being composed of both broad- and narrow-spiking waveforms, with some of the SOM^+^ cells co-localizing with excitatory neurons in the PhysMAP representations. This is entirely expected given the known heterogeneity of SOM^+^ cells which have variations in morphology as well as physiological properties relating to underlying sub-cellular types (Wu et al., 2023). In particular, many SOM^+^ cells demonstrate a fast-spiking phenotype in rodents and are known to express fast-spiking potassium channels (Urban-Ciecko and Barth, 2016). In contrast other SOM^+^ cells, such as Calb2, exhibit broad-spiking phenotypes (Hostetler et al., 2023). Similarly, in the juxtacellular dataset, excitatory neurons formed a broad class and layer 4 excitatory neurons had different and distinct electrophysiological properties from layer 5 excitatory neurons. While not directly investigated, this is in agreement with findings from intracellular recordings in motor cortex that corticothalamic- (deep layer 5 and layer 6) and intratelencephalic-projecting neurons (layer 2/3 and superficial layer 5) have distinct electrophysiological properties and morphologies (Scala et al., 2021). We anticipate this is at least in part why excitatory cells of different layers segregate in PhysMAP’s visualizations. These heterogeneities in SOM^+^ and excitatory cells, among others, are likely because cell types are not discrete and increasingly studies suggest they lie on a continuum. Said differently, variation within a discrete “cell type” is the norm and not the exception and can emerge from differences in morphology, laminar location, and gene expression (Cembrowski and Menon, 2018; Scala et al., 2021). This true underlying physiological variation is invisible to typical discrete classifier approaches but readily apparent with PhysMAP.

Second, PhysMAP offers a head-to-head comparison of each modality that many purely classifier methods do not provide easily. For instance, in both juxtacellular and extracellular recordings, waveform shape is the most reliable modality for delineating ground truth cell type. These results reaffirm our previous work (Lee et al., 2021) and with the observation that the expression of many genes which determine waveform shape often covary with neuronal cell types in different layers of cortex (Bomkamp et al., 2019). For instance, analysis of a mouse Patch-seq dataset revealed that the gene expression of many different potassium ion channels (e.g., *Kcnh7* and *Kcnc2*) correlated with properties of the waveform such as amplitude of after-hyperpolarization between excitatory and inhibitory cell types (e.g., Lamp5, Pvalb, etc, Bomkamp et al., 2019). Similarly, *Scn1b* is a gene encoding a voltage-gated sodium channel whose expression correlates with the width of the spike and many other electrophysiological properties (e.g., the first principal component) for both excitatory and inhibitory neuron classes. Thus, based on this data, it is perhaps expected that waveform shape would be one of the most reliable electrophysiological modalities for delineating cell types *in vivo*. In addition, this close covariation between transcriptomics and electrophysiological properties is perhaps the reason why our PhysMAP and previous WaveMAP approaches are successful in delineating transcriptomic cell types from electrophysiological recordings.

When conducting this study, we also realized that a key requirement for PhysMAP is high-quality ground truth datasets that cover the full range cell types. However, high-quality single unit recordings with opto-tagging are difficult to obtain for at least two main reasons: opto-tagging yield/specificity and single unit isolation.

First, opto-tagging experiments are technically challenging due to light attenuation through brain tissue or lack of transgene expression/specificity (Li et al., 2019), yield per recording is especially low when searching for less common cell types such as VIP^+^ cells which compose only *∼*13% of inhibitory neurons (Prönneke et al., 2015). These physically small cell types like VIP^+^ are often “missed” even on high-density probes. but are detected on *ultra* high-density probes possibly due to increased signal quality and denser sampling (Ye & Shelton et al. 2023). Furthermore, most opto-tagging protocols isolate directly tagged cells by examining the first 10 ms after the onset of a light pulse (Lima et al., 2009). However, Beau et al. (2024) find that some cells that fire within this 10 ms window after stimulation, have their spiking abolished by the introduction of blockers of synaptic transmission. Thus, more experiments are needed to refine opto-tagging experimental protocols and with devices that are able to catch all cell types in an unbiased manner.

Second, single unit isolation is a significant challenge for electrophysiology (Buccino et al., 2022). Even with modern recording technologies (Jun et al., 2017) and spike sorting methods (Pachitariu et al., 2016), single neuron isolation is difficult (Vincent and Economo, 2023). Poor isolation will lead to a multi-unit being classified as a single neuron and impacts PhysMAP because it blurs the physiological differences between cell types. For example, averaging together waveforms from a fast-firing narrow-spiking PV^+^ cell and a slow-firing broad-spiking excitatory pyramidal cell could lead to an artifactual intermediate width waveform neuron that is neither PV^+^ nor an excitatory pyramidal neuron. This mixing of cell types would complicate inferences from PhysMAP.

We anticipate that these technical challenges will be overcome. However, one key need for PhysMAP and other cell type classifiers is a more exhaustive characterization of the many cell types in the brain. Inhibitory cell types receive the most attention in opto-tagging experiments whereas excitatory populations have typically been under explored. In the datasets we analyzed here, only one (CellExplorer data from Buzsaki lab) contains positive identification of excitatory cells and this from an older study that precedes optogenetics (Henze et al., 2000). This is possibly the result of reliance on classical heuristics in the field that broad-spiking cells are excitatory neurons (McCormick et al., 1985). However, not all broad-spiking cells are excitatory (Wu et al., 2023) and not all excitatory cells are broad-spiking (Vigneswaran et al., 2011). This complex relationship between spike width and cell types is especially true in primates (Lemon et al., 2021). Another possible reason for this lack of attention to direct tagging of excitatory cells is that it can be hard to distinguish which cells are being directly stimulated due to the density of labeling and degree of post-synaptic secondary stimulation (Li et al., 2019). One way to label excitatory neurons more sparsely and specifically, is to use projection targeting approaches identifying neurons with long-range compared as opposed to local connections (Economo et al., 2018). Building full cell type classifiers (for cortex) will necessitate the opto-tagging of major projection classes most notably the three major excitatory types (intratelencephalic-, extratelencephalic-, and corticothalamic-projecting) and also two of the major remaining major inhibitory cell types (*Lamp5* and *Sncg*). Furthermore, it should be explored the extent to which electrophysiological signatures of these cell types are shared between brain areas. The conserved transcriptomic identity of many cell types across areas seems to suggest that this is the case at least for cortex (Yao et al., 2023).

Thus, PhysMAP and other cell type classifiers (Beau et al., 2024) would benefit from dedicated opto-tagging experiments that assess a wide of cell types, prioritizing single unit isolation, and with robust opto-tagging yields. Fortunately, two technological advances will help with these experiments. First, Neuropixels probes have dramatically increased the number of recording sites and capture the same neuron across multiple channels to provide a “multi-channel waveform”. This has been shown to better capture morphological details which might be important for differentiating cell types that only differ in their structure (Jia et al., 2019; Ye et al., 2023; Sibille et al., 2022; Carr et al., 2024). Second, new enhancer-based viruses demonstrate excellent cell subtype-specificity (Green et al., 2023) along with efficacy across diverse organisms including humans (Graybuck et al., 2021). The combination of these technical advances with our PhysMAP approach will likely provide robust cell type classifiers that will allow the simultaneous assessment of the dynamics of neuronal cell types *in vivo* during sensation, perception, and action in multiple species.

## Methods

The methods are organized as follows. We first describe the datasets that we used for the paper and then describe the weighted nearest neighbor method from PhysMAP. Finally, we describe the various analyses used for quantifying the performance of PhysMAP relative to individual modalities and also other controls.

### Open Datasets

Here, we only briefly detail the most relevant aspects of each dataset for our purpose of validating our PhysMAP approach (Table 1). We refer the reader to each dataset’s respective publication for additional methodological information.

**Table 1:** The three datasets analyzed with the modalities used and the cell types identified in each.

| Study | N | Available modalities | Classifiable cell types | Data URL |
| --- | --- | --- | --- | --- |
| Yu et al. (2019) | 246 | Waveform shape, ISI dist., PSTH | PV<sup>+</sup>-4, PV<sup>+</sup>-5, SOM<sup>+</sup>, E-4, E-5 | Link |
| Lakunina et al. (2020) | 373 | Waveform shape, ISI dist. | PV<sup>+</sup>, SOM<sup>+</sup> | Link |
| Petersen et al. (2021) | 430 | Waveform shape, ISI dist., Auto-correlation, Derived ephys. metrics | PV<sup>+</sup>, SOM<sup>+</sup>, E | Link |

### Juxtacellular Mouse S1 Dataset

The juxtacellular dataset used in Fig. 2 and Fig. 3 and analyzed in section were collected by Yu et al. (2019) and downloaded from the associated file sharing portal (see Table 1). Recordings were performed in the primary somatosensory (barrel) cortex of Sst-IRES-Cre × Ai32, Pvalb-IRES-Cre/Pvalb-Cre × Ai32, or Vip-IRES-Cre × Ai32 mice. These experiments focused on *in vivo* juxtasomatic electrophysiological recordings using glass micropipettes. After the end of recording, the cells were filled with biocytin/neurobiotin. Filled neurons both acted as a verification of the recorded cell type via their morphology and also served as landmarks to align recording depths of each cell with cortical layers as determined by histology.

ChR2-expressing neurons were located during recording by observing laser-evoked spikes that occurred 1-2 ms after the onset of brief light pulses (5 ms) delivered at low frequency (5 Hz) for 3 out of every 10 s. Spikes were examined by eye and non-spike events were removed after projection in PCA-space. To control for drift, only waveforms between the 25th and 75th percentile of all spikes from a cell were used in further waveform shape analyses (average waveform and waveform shape metrics). Averaged waveform shapes and ISI distribution were extracted for each neuron from the *spkwaveform_all*.*mat* file in the corresponding cell type structs. In addition, the waveform shape metrics of spike width and peak-to-trough ratio were also collected from this file. Cells labeled “fake SOM^+^” were reassigned to the PV^+^population and cells labeled “putative SOM^+^” were reassigned to SOM^+^. This reassignment was based on a careful investigation in Yu et al. (2019) confirming earlier findings that off-target recombination occurs in fast-spiking neurons of somatostatin-IRES-Cre mice possibly due to transient Cre expression during development (Ma et al., 2006; Hu et al., 2013). PSTHs were calculated only from whisker touches during active whisking and activity resulting from both whisker protraction and retraction were used. Spiking was aligned to touch-onset and a PSTH calculated using 1 ms time bins. As not all cells contained PSTHs, only the subset of units containing all three modalities (waveform shape, ISI distribution, and PSTH) were used. To create the “concatenated” modality, we simply concatenated the waveform shape, ISI distribution, and PSTH for a given unit into a single long feature vector. Waveform metrics—peak-to-trough duration and peak-to-trough ratio—were calculated in the originating publication as the time from the first peak to the first trough and the ratio of absolute values of the peak to the trough respectively.

### Extracellular Mouse A1 Dataset

The extracellular data we analyzed from Lakunina et al. (2020) was downloaded from GitHub (see Table 1). PV^+^and SOM^+^ cells were identified by opto-tagging in Pvalb-Cre or Sst-Cre mice respectively that were the progeny of crossing LSL-ChR2 mice. Photoidentification of ChR2-expressing cells was relatively conservative with positively identified cells being those that exhibited significant firing rate changes (p < 0.001) in firing rate during the first 10 ms of stimulation-onset. Recordings were collected from primary auditory cortex via silicon probe and stimulation was via an optical fiber located at the top of the probe recording sites. Spikes were sorted offline in Klustakwik and the units used in further analyses only if ISI violation rate (number of spike intervals < 2 ms) was less than 2% per cluster. In the original publication, only cells with spike quality index (SQI; ratio between the peak amplitude of the waveform and the average variance, calculated using the channel with the largest amplitude) above 2.5 were used; in our analysis, we only used cells with SQI above 4.0. Mean waveforms were calculated for a unit using the channel with greatest amplitude of spikes.

### Extracellular Mouse Visual Cortex and Hippocampus Dataset (CellExplorer)

The third dataset was obtained from the CellExplorer package (Petersen et al., 2021) and is heterogeneous being composed of two separate extracellular electrophysiological data from the Allen Institute’s Brain Observatory and the Buzsaki lab and included waveform shape, ISI distribution, autocorrelogram (ACG), and various derived electrophysiological metrics. We used the derived eletrophysiological metrics that all units shared and this included the following eleven:

- Spike width (*troughToPeak*)
- Peak-to-trough derivative (*troughtoPeakDerivative*)
- Pre-hyperpolarization peak-to-post depolarization peak ratio (*ab_ratio*)
- Coefficient of variation (*cv2*) measuring ISI distribution
- ACG tau rise (*acg_tau_rise*)
- ACG tau decay (*acg_tau_decay*)
- ACG tau bursts (*acg_tau_burst*)
- ACG refractory period (*acg_refrac*)
- ACG decay amplitude (*acg_c*)
- ACG rise amplitude (*acg_d*)
- ACG burst amplitude (*acg_h*).

More information on these metrics are available on the CellExplorer website docs (link). For the analyses, these metrics were concatenated for each neuron into a single vector per-unit to form the electrophysiological metrics modality. Given the large number of publications and protocols that have contributed to the CellExplorer dataset, we only provide a brief overview here.

### CellExplorer: Allen Institute Data

Data from the Allen Institute was collected from mouse primary visual cortex (V1) and higher visual areas (HVAs) using simultaneous Neuropixels probe recordings (Siegle et al., 2021). Opto-tagging was conducted for PV^+^, SOM^+^, and VIP^+^ cell types using Pvalb-IRES-Cre × Ai32, Sst-IRES-Cre × Ai32, and Vip-IRES-Cre × Ai32 mice respectively. Stimulation was conducted at the end of each behavior session via blue LED/laser through an optical fiber with surface illumination of the cranial window using a rounded 10 ms square pulse. Tagged cells were identified by an average two times increase in base firing rate during stimulation relative to baseline.

### CellExplorer: Buzsaki Lab Data

Data from the Buzsaki lab were collected under several different protocols but CA1 excitatory neurons were identified by paired intra- and extra-cellular recordings (Henze et al., 2000). V1 PV^+^cells from PV-Cre × Ai32 mice were identified as those that were stimulated to over 8 standard deviations of the mean of baseline in the first 6 ms of stimulation with a 10 ms square pulse (Senzai et al., 2019). Hippocampal PV^+^cells were obtained from PV-Cre × Ai32 using a 50 -100 ms light pulse and identified as tagged if they 1) had a statistically significant increase in firing rate and 2) if the stimulated firing rate was over 50% of the baseline (English & McKenzie et al. 2017).

### PhysMAP Preprocessing

Modalities from all neurons were preprocessed via the same steps required by and performed using Seurat v4:

1. **PCA reduction**: All modalities were initially dimensionality reduced to the smallest input modality dimension. For the S1 dataset, this is 30-dimensions; for the A1 dataset, this is 20-dimensions; and for CellExplorer, this is 40-dimensions.
2. **Normalization**: Each modality is separately normalized using a centered log ratio transform.
3. **Rescaling and centering**: Each feature of each modality is linearly rescaled to occupy the same unit variance. Each feature is then centered by mean subtraction.

### Weighted Nearest Neighbor (WNN) Algorithm

The PhysMAP approach uses the WNN algorithm available from Seurat v4 (Hao & Hao et al. 2021) which is summarized as the following steps.

1. **Nearest neighbor identification**: Within the preprocessed, high-dimensional space of a given modality (here, waveform shape Fig. 1A), a single unit is selected (Fig. 1B, blue sphere) and its nearest neighbors identified via Euclidean distance in the ambient (un-dimensionality reduced) space (Fig. 1B, red spheres).
2. **Nearest neighbor prediction (within modality)**: A prediction of the selected single unit’s waveform is made via unweighted averaging the *k*-nearest neighbors identified in Fig. 1B. By default, we use *k* = 20 and Euclidean distances in the ambient space to locate the nearest neighbors.
3. **Within-modal affinity calculation**: The unit’s nearest neighbor predicted waveform shape generated by *k*-nearest neighbors 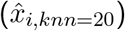 and its actual waveform shape (*x*_*i*_) are then passed into a modified UMAP distance kernel,

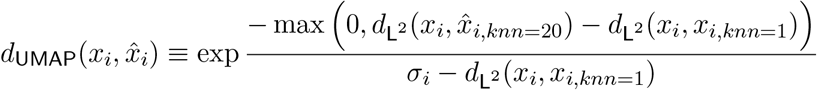

where 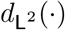 is the Euclidean (L^2^-norm) distance metric; 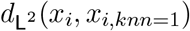 is the Euclidean distance from *x*_*i*_ to its nearest neighbor; and *σ*_*i*_ is a modified UMAP bandwidth equal to the average of the Euclidean distances from the *i*^th^ unit to the 20 nearest units with lowest non-zero Jaccard index. This equation is used to calculate the “within-modal affinity” which gives a measure of how predictive a certain modality is of this unit. This forms the numerator of the “affinity ratio” shown in Fig. 1D.
4. **Nearest neighbor identification (cross modality)**: Examining normalized single unit ISI distributions (the “cross-modal” space) generated from the same neurons in Fig. 1E, we locate the same unit and nearest neighbors previously identified but in ISI distribution-space Fig. 1F.
5. **Nearest neighbor prediction (cross modality)**: As in Fig. 1C, a nearest neighbor prediction is made but this time, in a different modality albeit using the same neurons identified in the original modality Fig. 1G, red dashed circle.
6. **Affinity ratio calculation**: This unit and its prediction in the cross modality (ISI distribution-space) are used to compute the “cross-modal affinity” by passing it into a modified UMAP distance kernel; this forms the denominator of the waveform affinity ratio for a given unit (*S*_Wave_(*i*)) shown in Fig. 1D. This process is repeated for every unit in the dataset and also in the reverse manner: beginning with the ISI distribution-space as the within modality and with the waveform shape-space as the cross modality. For datasets with more than two modalities, the affinity ratio is calculated for each modality versus every other and for every unit. Thus for a dataset with *d* data points and *n* feature modalities, there will be 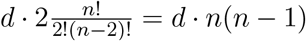 affinity ratios to calculate.
7. **Converting affinity ratios to modality weights**: With affinity ratios calculated for both waveform and ISI distribution across all units, modality weights (*β*_Wave_(*i*) and *β*_ISI_(*i*)) for each unit can be calculated. For a given unit, this weight is the ratio of a given modality’s affinity ratio for that unit divided by the sum of all other modality’s affinity ratios for that unit. For the two-modality case, this is summarized by the equation in Fig. 1H.
8. **Calculating unit pair-wise affinities**: For every pair of units, an affinity is also calculated by passing both of them and their coordinates into UMAP’s distance kernel. This occurs for the same pair of units in both the waveform shape-space Fig. 1I, top and the ISI distribution-space Fig. 1I, bottom to determine pair-wise affinities *θ*_Wave_(*i, j*) and *θ*_ISI_(*i, j*) respectively. This is done for all pairs of points and all modalities.
9. **Creating the WNN**: To create the weighted-nearest neighbors representation, a modality-weighted sum of the pair-wise affinities Fig. 1J, top is taken to produce a “connectivity matrix” Fig. 1J, bottom. The *k*-nearest neighbor algorithm is applied to this matrix to form the final WNN (default number of neighbors is *k* = 200) which is then visualized into two dimensions with UMAP’s force-directed graph layout projection for visualization.

### Classification Analysis

After a WNN is constructed for a dataset, we used UMAP’s force-directed graph layout procedure to project the graph into a 10-dimensional embedded space. We then used a random 15-85% test-train split and trained a stochastic gradient boosted tree model (GBM) classifier on the training set with five-fold cross-validation using the *caret* package in R (Kuhn, 2008) to identify each underlying cell type with multiclass (one-vs-rest) objective function. We then repeated this train-test split 20 times and averaged the balanced accuracy over these independent runs. Each classifier also underwent hyperparameter tuning via a grid search. This procedure was run identically for the controls in Fig. S2A, B except with varying graph embedding dimension and using different classifiers available in the *caret* package.

### Leiden Community Detection

To calculate the MARI score for each dataset across different numbers of clusters we used Leiden clustering (Traag et al., 2019) on each dataset’s WNN graphs with resolution between 0.1 and 3.0 in 0.1 resolution intervals. The Leiden algorithm is a method for “community detection” which finds highly inter-connected nodes on a network graph akin to clustering in a metric space. Algorithms like Leiden (and the simpler Louvain algorithm) attempt to find a partitioning of the network into a set of communities that maximizes the modularity of each community. This modularity, *ℋ*, is a measure of how inter-connected the nodes are within a community versus outside of it. In the below definition, *m* is the total number of edges in the network, *e*_*c*_ is the number of edges in a community *c, γ* is the resolution parameter, and *K*_*c*_ is the sum of the degrees of the nodes in community *c*.

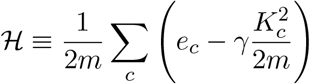

Thus, the resolution parameter is effectively a prior on the number of expected communities that should be found during the optimization; with lower resolution, less communities are found and with higher resolution, more communities are found.

### Modified Adjusted Rand Index (MARI) Calculation

Once this partitioning of a graph into communities is computed, a MARI score (Sundqvist et al., 2020) is calculated between each of these each neuron’s community membership and the ground truth cluster identity to determine how closely they corresponded. To understand MARI, which is a modification of the adjusted Rand index (ARI), we begin with an explanation of the Rand index. Given two clusterings, *X* and *Y* of the same dataset, the number of pair-wise elements that share cluster membership in both *X* and *Y* is *a* and the number of pair-wise elements that share different cluster memberships is *b*. The Rand index (*RI*) is 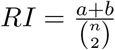 where *n* is the total number of pairs. The Rand index can also be interpreted as the sum of the number of true positive and true negative correspondences divided by the total number of guesses if one clustering is regarded as a classifier prediction of the other. However, in the classifier interpretation, the Rand index is not corrected for chance predictions. The ARI corrects for these chance predictions by subtracting the expected Rand index under the hypergeomtric distribution given two independent clusterings and fixed number of clusters in each. MARI further refines this metric by instead incorporating a multinomial distribution which does not enforce cluster size and is a better assumption given that most clustering algorithms (including Leiden) do not fix cluster sizes. This was used to compare each of the Leiden clusterings at different resolutions to underlying ground truth clusters.

### Reference Mapping with Anchors and Cell Type Classification

Unlike the previous classification analyses, an 80-20 train-test split was conducted before applying PhysMAP. This was done to combat bias in cross-validation because of data leakage due to pre-processing transformations (Moscovich and Rosset, 2022). The training set was used to create the reference mapping upon and the test set was used to create the query dataset. To construct the reference mapping, an WNN was constructed from each neuron’s waveform shape, ISI distribution, autocorrelogram, and derived electrophysiological metrics as before. Following the procedure for multi-modal reference mapping recommended in Hao & Hao et al. 2021, we first ran supervised principal component analysis (SPCA) on the reference data. Next, the query dataset is projected onto the reference using the previously computed SPCA transform. Now that reference and query datasets occupy the same space, “anchors” can be calculated between the reference and query. These anchors are pairs of cells between the reference and query that are located within each other’s neighborhoods; this concept is also referred to as mutual nearest neighbors (MNN). These neighborhoods are defined by computing *k*-nearest neighbors with *k* = 5. This nearest neighbor search is conducted in only the top-30 SPC’s. Once all anchors are found, they are scored which is an assessment of how confident we are in their correspondences. This score is calculated for each anchor given the 30 nearest neighbors in each reference and query dataset for both the reference and query data points of the anchor respectively. The overlap of these nearest neighbor matrices between the 0.01 and 0.90 quantiles is then linearly rescaled to be between 0 and 1 for all reference-query anchor pairs; this provides a score for each anchor. A weight matrix is then constructed between each query cell *c* and each *i*th nearest anchor *a*_*i*_ (where *i* ∈ [1, 50]) based upon the distance to each query cell and the corresponding anchor score. These weighted distances *D*_*c,i*_ are calculated as

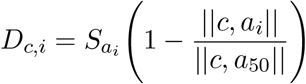

where 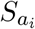 is the score of the *i*th anchor where || *·* || is the Euclidean distance. These distances are passed through a Gaussian distance kernel 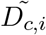 as,

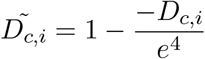

to form the entries of a weight matrix

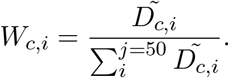

For cell type classification, a classification matrix *L* is created where each row corresponds to a ground truth cell type class and each column corresponds to each reference anchor. If a certain reference anchor pertains to a certain cell type, it is given a 1 and a 0 if otherwise. Label predictions *P* are then computed simply as,

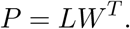

This returns a prediction score for each cell in the query dataset. A query cell is thus given a predicted cell type of whichever class has the highest prediction score.

## Author Contributions

EKL and CC jointly developed the idea of a cell type classifier and ultimately a lookup table based on ideas from Dr. Karel Svoboda, and Dr. Josh Siegle. AG helped with derived waveform metric analysis and several control analyses. GH helped collect one of the open datasets and provided technical advice especially with regards to the usage of quality metric and helped edit initial manuscript drafts. AL and SJ provided technical support and also collected one of the open datasets. PFP provided guidance on various approaches for multi-modal integration and other technical advice. Both EKL and CC curated datasets. CC wrote initial R code and performed analyses. Code was further refined and augmented by EKL. EKL made figures and wrote initial drafts of the paper with CC.

## Acknowledgments

We are grateful to Dr. Karel Svoboda, Dr. Anna Lakunina, and Dr. Josh Siegle for the initial ideas of cell type classifiers and lookup tables for electrophysiology. We also thank Dr. Yujin Han, Dr. Josh Siegle, Pierre Boucher, Nicole Carr, Tian Wang, Vivan Moosmann, Tushar Arora, and Munib Hasnain for their thoughtful comments on our manuscript.

