## Supplemental Figures for "PhysMAP - interpretable *in vivo* neuronal cell type identification using multi-modal analysis of electrophysiological data"

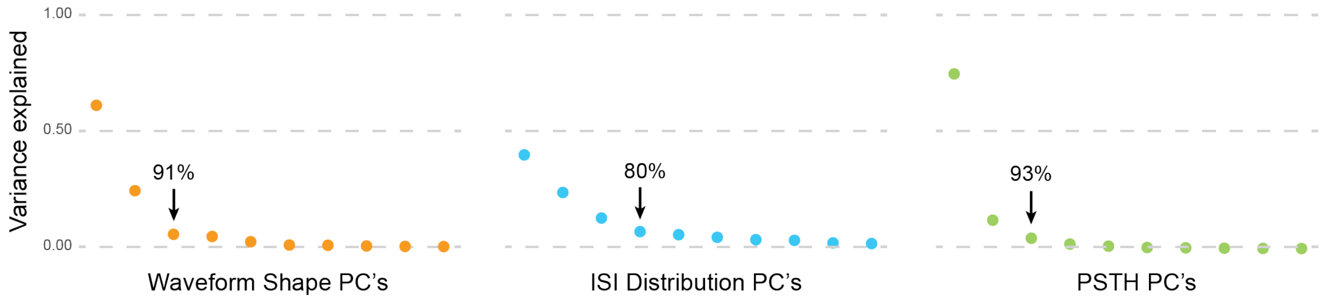


Figure S1: **Principal component analysis gives an estimate of linear intrinsic dimensionality**: Principal component analysis (PCA) was applied to the data from each modality in Fig. 2A and the per-component variance explained calculated for each. For each modality, the scree plot shows the variance explained for each principal component. An arrow on each plot demarcates the "elbow" with cumulative percent variance explained listed above.

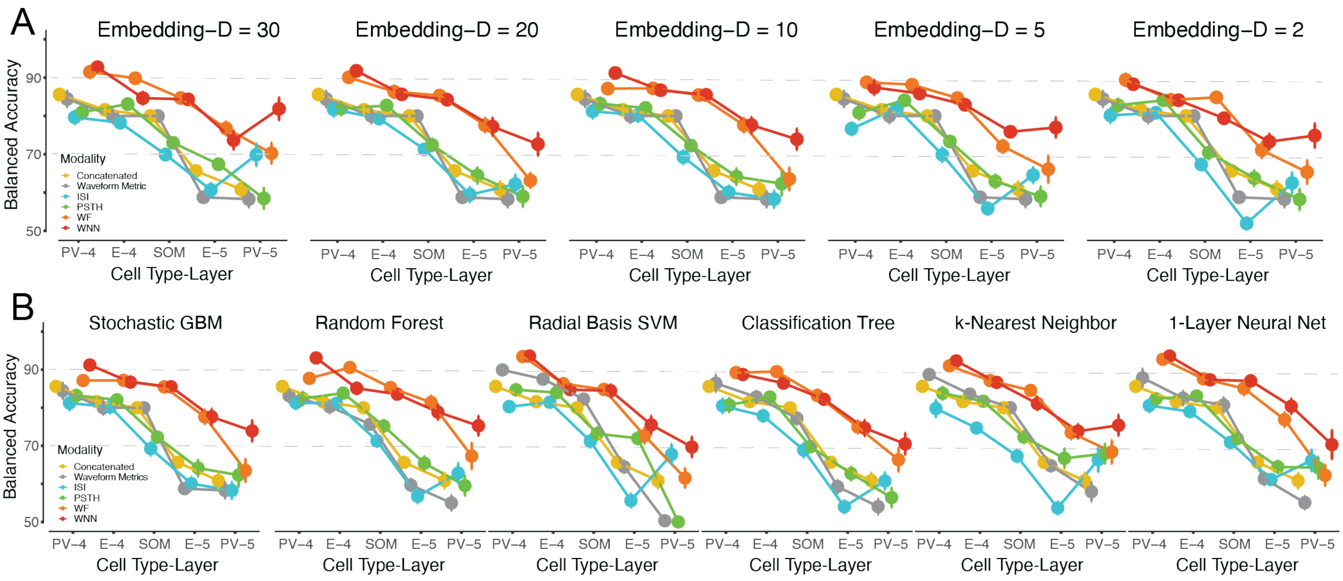


Figure S2: **Classifier performance is not affected by graph embedding dimensionality or choice of classifier**: (**A**) The embedding dimensionality of the WNN graph was varied between 30 and 2 dimensions and used for training the same GBM classifier as in Fig. 3A which was trained on embedding dimension of 10 with five-fold cross-validation. The mean balanced accuracy (± S.E.) across the five cell types with more than 25 examples is shown. (**B**) With a WNN graph embedding dimension of 10, six different classifiers (with default *caret* settings) were trained to identify the five cell types in Fig. 3A (which was the same here as the stochastic GBM). These plots are once again shown with mean balanced accuracy (± S.E.) after five-fold cross-validation.

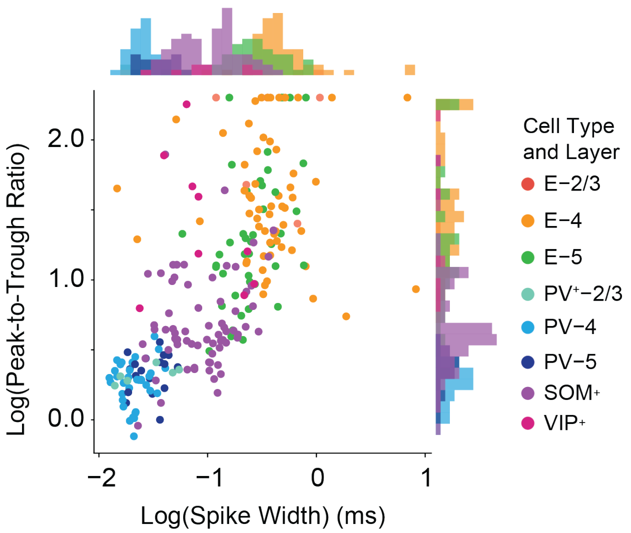


Figure S3: **Waveform shape metrics offer intuition but are disconnected from PhysMAP**: The neurons in Fig. 2 now in a scatter plot according to the log transformation of spiked width (in ms) and peak-to-trough ratio. Histograms for each metric are shown on the opposite marginals. This is a reproduction of the plot in Fig. 1E of Jia et al. (2019).

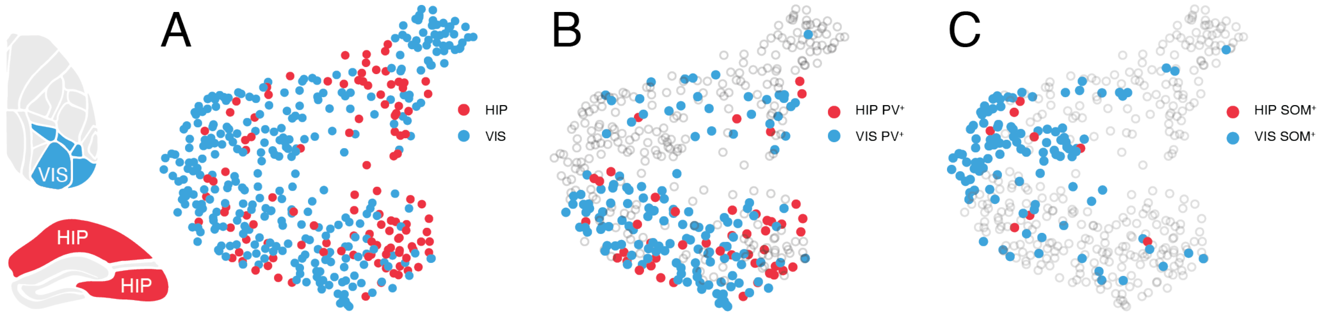


Figure S4: **Cell types are identified similarly between brain areas**: (**A**) Neurons in PhysMAP colored by their origin in hippocampus (red) or visual cortex (blue) of the mouse in the CellExplorer dataset (Petersen et al., 2021). (**B**) The same neurons in (A) but now restricted only to PV$^+$ cells. (**C**) The same neurons as in (A) but now restricted to only SOM$^+$ cells.
